# Epidermal tyrosine catabolism is critical for metabolic homeostasis and survival against high-protein diets in *Drosophila*

**DOI:** 10.1101/2023.04.20.537645

**Authors:** Hina Kosakamoto, Masayuki Miura, Fumiaki Obata

## Abstract

The insect epidermis that forms the exoskeleton and determines the body size of an organism has the potential to respond and adapt to the changing nutritional environment. However, the extent to which the tissue contributes to adaptation to varying dietary protein availability, as well as its role as a metabolic regulator, remains incompletely understood. Here, we show that the *Drosophila* epidermis promptly reacts to dietary protein intake, promoting tyrosine (Tyr) catabolism. Among the enzymes involved in Tyr degradation, 4- hydroxyphenylpyruvate dioxygenase (Hpd) is particularly induced under high-protein regimens. We found that AMP-activated protein kinase (AMPK) pathway and forkhead box O subfamily (FoxO) induce Hpd expression. Impaired Hpd function in the epidermis leads to aberrant increases in internal Tyr levels and its metabolites, disrupting larval development under high-protein diets. Taken together, our findings uncover the critical role of epidermal metabolism in adapting to imbalanced protein environments and hence in promoting animal survival.

**Summary statement:** Tyrosine degradation is upregulated in *Drosophila* epidermal tissue via the AMPK-FoxO axis upon dietary high-protein stress.

## Introduction

Metabolic adaptation to an altered nutritional environment is pivotal for animals. Since both shortage and excessive nutrition can be detrimental, animals are equipped with various mechanisms to maintain metabolic homeostasis. *Drosophila melanogaster*, a model organism with evolutionarily conserved metabolic pathways and its capability for efficient genetic manipulation, has provided valuable insights into the mechanisms of metabolic adaptation (Chatterjee and Perrimon, 2021). Among a myriad of nutrients, dietary protein, which serves as a precursor to amino acids (AAs), exerts a profound influence on animal physiology including the growth, fertility, and lifespan of fruit flies (Soultoukis and Partridge, 2016). Flies can modulate metabolism to maintain organismal homeostasis by detecting and adapting to alterations in dietary protein availability. Multiple organs are involved in AA perception. Primarily, sensory organs or neurons located in the gut rumen perceive the taste of AAs, which then transmit the signal to the central nervous system (Croset et al., 2016; Steck et al., 2018; Toshima et al., 2012; Yang et al., 2018).

Subsequently, digested AAs can be directly sensed in the brain, eliciting neurotransmission (Bjordal et al., 2014; Manière et al., 2016), or in peripheral tissues, which often involves the secretion of hormones, triggering systemic adaptive responses (Agrawal et al., 2016; Delanoue et al., 2016; Koyama and Mirth, 2016). The fat body represents a peripheral organ that plays a crucial role in AA sensing (Li et al., 2019). In response to a shortage of dietary amino acid intake, the deactivation of mechanistic target of rapamycin complex 1 (mTORC1) signalling in the fat body affects several endocrine hormones including Stunted, Growth-blocking peptide 1 and 2, and Eiger (Manière et al., 2020). These fat body-derived soluble factors regulate the secretion of *Drosophila insulin-like peptides* (*Dilps*) from the brain that systemically stimulate insulin-like receptor (InR) to activate the insulin/IGF-1- like signalling (IIS) pathway. IIS governs both growth and metabolism, partly through its downstream transcription factor FoxO, which targets various metabolic enzymes (Chatterjee and Perrimon, 2021; Link and Fernandez-Marcos, 2017). The fat body is believed to be a major metabolic organ, functioning as a counterpart of mammalian liver and white adipose tissue where many metabolic pathways are active. However, the contribution of other peripheral organs to maintaining metabolic homeostasis in response to changing protein environments is poorly understood.

Especially in insects that possess an exoskeleton, nutritional sensing in the epithelial tissue is supposed to be of great importance, because it limits the animal’s body size. Indeed, epidermal cells exhibit a highly responsive nature to genetically altered nutritional signalling, such as insulin receptor/phosphoinositide 3-kinase (PI3K) signalling, resulting in a marked alteration of cell size (Britton et al., 2002). Since the epithelium covers the entire body, it is fully exposed to haemolymph and possesses a great capacity to modulate circulating metabolites. Despite the long-standing examination of the epidermis’s role in barrier function and cuticle formation, which includes sclerotization and tanning during metamorphosis (Hopkins and Kramer, 1992), the extent to which the tissue responds and adapts to varying dietary protein availability, as well as its role as a metabolic regulator within the organism, remains to be elucidated.

Tyrosine is a nonessential amino acid critical for catecholamine and melanin synthesis in addition to protein synthesis. Previously, we revealed that tyrosine (Tyr) plays a crucial role in sensing dietary protein sufficiency (Kosakamoto et al., 2022). The scarcity of Tyr is perceived through activating transcription factor 4 (ATF4) in the fat body, which alters protein synthesis, mTORC1 signalling, and feeding behaviour. In contrast, excessive accumulation of Tyr can be detrimental due to its limited solubility and propensity to precipitate within cells as crystals (Scott, 2006; Wu, 2013). Hence, the metabolic homeostasis of Tyr during excessive protein consumption should be tightly regulated.

Internal levels of Tyr can be modulated through its biosynthesis and degradation processes. Tyr is derived from phenylalanine (Phe), with approximately 70% of it continuously undergoing degradation into fumarate and acetoacetate for energy utilization (Shiman and Gray, 1998). However, the contribution of Tyr catabolism to metabolic adaptation and its regulatory mechanisms upon protein feeding remain unresolved.

In this study, we have unveiled the intricate regulation of enzymes responsible for the degradation of Tyr, which are conspicuously expressed in the epidermis. Their expression is highly responsive to dietary protein levels. The upregulation of Tyr degradation enzymes is imperative for metabolic homeostasis and development under high-protein diets, thereby underscoring the epidermis as a metabolic regulator during altered nutritional input.

## Results

### Tyrosine metabolism is upregulated by a high-protein diet in the larval body wall

To investigate the metabolic adaptation of the epidermis to changes in the supply of dietary protein, we conducted RNA sequencing analysis of the larval body wall, which comprises mainly the epidermis but also contains muscle, haemocytes, neurons, and oenocytes. To manipulate the dietary protein composition, we fed third instar larvae diets consisting of yeast extract (YE), the concentration of which varied among 4% (low), 10% (normal), and 24% (high). Feeding these diets for six hours changed the gene expression profile in the body wall (Figs. 1A-D). Gene Ontology analysis revealed that the genes responsible for protein targeting from the endoplasmic reticulum to the Golgi apparatus and chaperones were induced by feeding the high-protein diet (Fig. 1C). Additionally, Phe and Tyr catabolic processes were significantly upregulated (Fig. 1C, Table S1). These results suggest an acceleration of protein processing and AA degradation in response to the high load of dietary protein. Concurrently, the expression of AA transporters and glutamine synthetase 2 was suppressed (Fig. 1D), indicating that the import and biosynthesis of AAs are downregulated to maintain AA homeostasis.

**Fig. 1.**
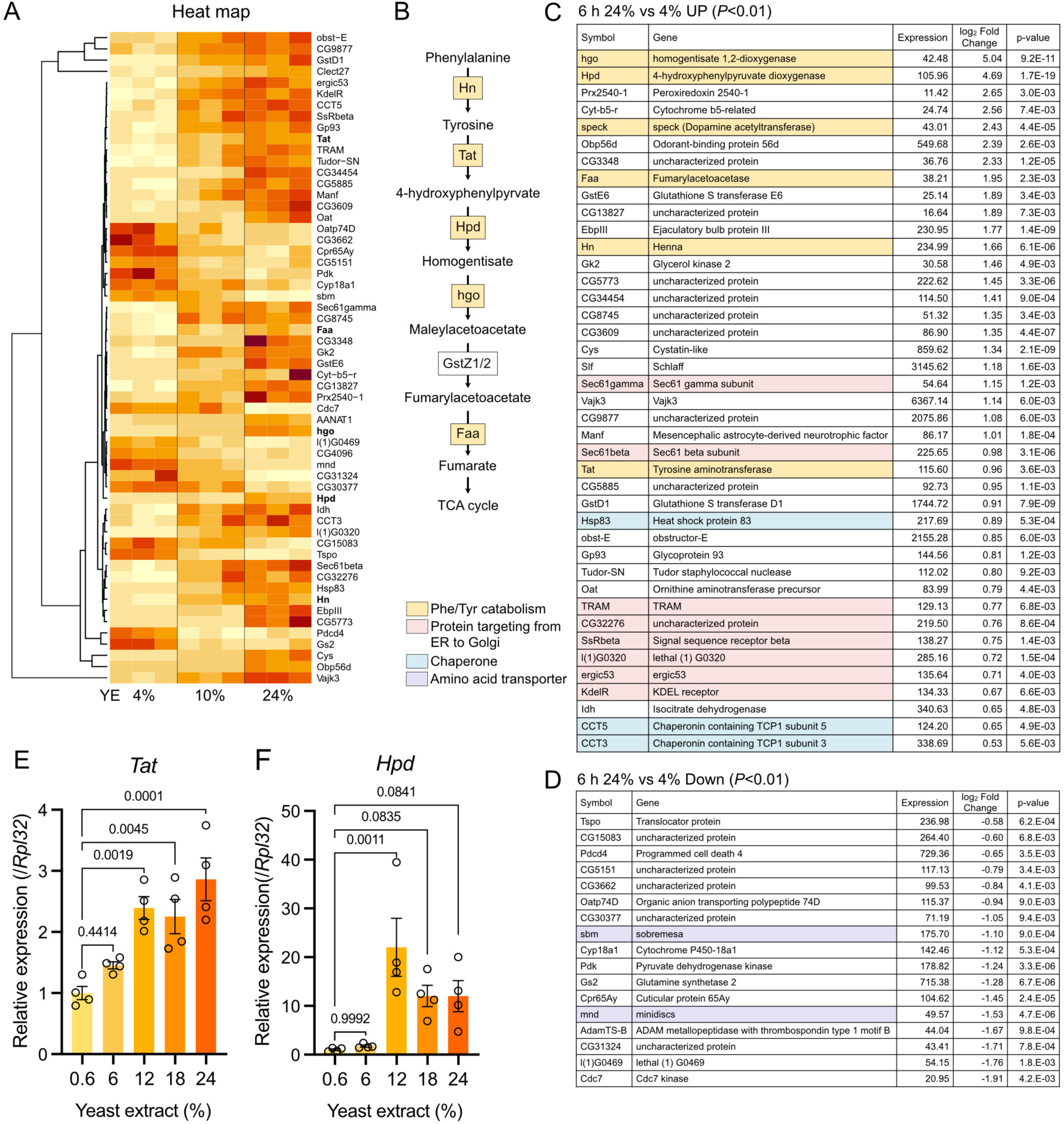
Tyrosine catabolism is increased in response to high-protein feeding. (A) Heatmap analysis of differentially expressed genes (DEGs) from the RNAseq analysis of the carcass of *w^Dah^* larvae upon a high-protein diet. n = 3. (B) Tyr degradation pathway. Genes marked with orange were induced upon a high-protein diet. (C, D) Lists of up- (C) and down- (D) regulated genes in the larval carcass upon a high-protein diet. (E, F) Quantitative RT‒PCR of *Tat* (E) and *Hpd* (F) in the whole body of third instar larvae fed different protein diets for eight hours. n = 4. For all graphs, the mean and SEM with all data points of biological replicates are shown. P-values were determined by one-way ANOVA with Dunnett’s multiple comparison test.

It was intriguing that the Phe/Tyr degradation pathway was specifically induced when high protein intake led to an increase in AA levels in general. To determine the threshold and dose‒response relationship of the regulation of Phe/Tyr degradation enzymes, we performed quantitative RT‒PCR for the first and second enzymes in the tyrosine degradation pathway, *tyrosine aminotransferase* (Tat) and *4-hydroxyphenylpyruvate dioxygenase* (Hpd), respectively. Both genes were induced between 6% and 12% YE concentrations, although there was a tendency towards a gradual increase, especially for *Tat* (Figs. 1E, F).

### Hpd is expressed in epidermal cells

The remarkable increase in *Hpd* transcripts upon a high-protein diet motivated us to perform a detailed analysis of its molecular regulation. We utilized the CRISPR/Cas9 system to introduce monomeric ultrastable GFP (muGFP, (Scott et al., 2018)) at the C-terminus of endogenous *Hpd*. As anticipated, the GFP fluorescence of *Hpd::muGFP* larvae increased in response to higher YE concentrations (Figs. 2A, B). Notably, the expression was limited to the body wall, especially on the dorsal side, and was not observed in visceral metabolic organs such as the fat body or the gut (Fig. S1A). This finding is consistent with the expression pattern demonstrated in the FlyAtlas Anatomy Microarray (Fig. S1B, Flybase.org). Not only *Hpd* but also other Tyr catabolic enzymes are highly expressed in the carcass, although *Henna* (*Hn*) is also highly expressed in the fat body (Fig. S1B, Flybase.org). Combining *Hpd::muGFP* with *UAS-mCD8-RFP* driven by the epidermis driver *A58-Gal4*, we discovered that Hpd reporter expression merged with epidermis-driven RFP and not with the muscle under the epidermis stained by phalloidin (Fig. 2C). Magnified images showed that Hpd::muGFP was localized to the cytosol and nucleus (Fig. 2D). This result suggests that Tyr may be degraded in the nucleus as well as the cytoplasm, although we cannot deny the possibility that muGFP fusion perturbs protein localization. In adult flies, we also observed an increase in Hpd expression upon higher protein consumption, and the expression was restricted to the epidermis of the head and thorax and oenocytes (Fig. 2E, Fig. S1C).

**Fig. 2.**
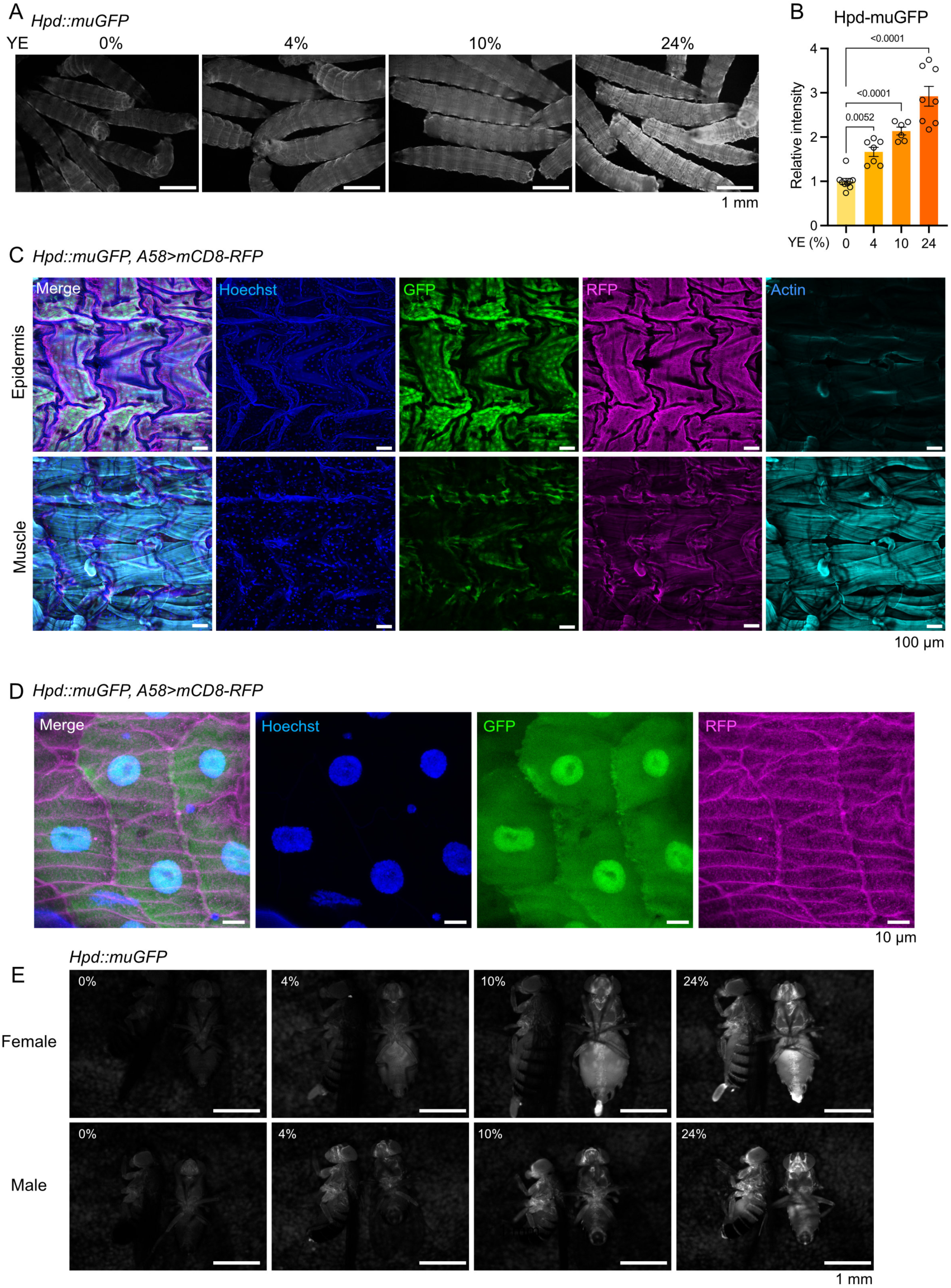
Hpd is expressed in the epidermis. (A, B) Images (A) and quantification (B) of *Hpd::muGFP* larvae upon various concentrations of dietary protein. Scale bar: 1 mm. n = 6-9. For the graph, the mean and SEM with all data points of biological replicates are shown. P-values were determined by one-way ANOVA with Dunnett’s multiple comparison test. (C, D) Images of dissected body walls of *Hpd::muGFP, A58>mCD8-RFP* larvae. Nuclei are stained with Hoechst 33342. Actin in the muscle layer is stained with Phalloidin-iFluor 647. Scale bar: 100 µm (C) or 10 µm (D). (E) Images of lateral and ventral views of *Hpd::muGFP* adults upon various concentrations of dietary protein. Scale bar: 1 mm

Although *A58-Gal4* is often used for genetic manipulation in the epidermis, its expression has also been observed in other tissues, including the fat body, gut, Malpighian tubules, and salivary gland. Since Hpd expression is highly specific to the epidermis, we generated a GeneSwitch driver using the 500 bp upstream region of the *Hpd* start codon for drug-inducible gene manipulation in the epidermis. When the driver line was crossed with *UAS-GFP* flies, GFP was visible in the epidermis only when mifepristone (RU486), a drug that activates GeneSwitch, was added to the food (Fig. S2A). As expected, Hpd expression in the epidermis was suppressed by *Hpd* knockdown using the driver with RU486, suggesting the feasibility of this tool for epidermis-specific gene manipulation (Fig. S2B).

Notably, it has been shown that the Tyr degradation pathway is upregulated during ageing (Parkhitko et al., 2020). Amelioration of degradation by suppression of *Hpd* can extend organismal lifespan (Parkhitko et al., 2020). Consistently, we also found that the Hpd::muGFP reporter fluorescence in adult females increased during ageing, especially at three weeks of age, while males showed little change (Fig. S2C). Therefore, the regulation of Tyr degradation in the epidermis may be relevant for ageing.

### Hpd induction is mediated by FoxO in response to Tyr feeding

We next asked how Hpd was induced by protein intake. As it is responsible for catalysing Tyr degradation, sensing the quantity of Tyr should be critical in the regulation of *Hpd* expression. We found that supplementing a high amount of Tyr alone to a standard yeast-based diet induces Hpd expression, whereas neither the other nine nonessential AAs (NEAAs) nor the nine essential AAs except for Phe had an effect (Figs. 3A, B). Decreased Hpd expression upon a low protein diet was recovered by adding back Tyr (Fig. 3C). Additionally, specific Tyr restriction for three days suppressed Hpd expression in adult females, while males showed minor effects (Fig. S3A), suggesting that dietary and/or internal Tyr levels modulate Hpd expression.

**Fig. 3.**
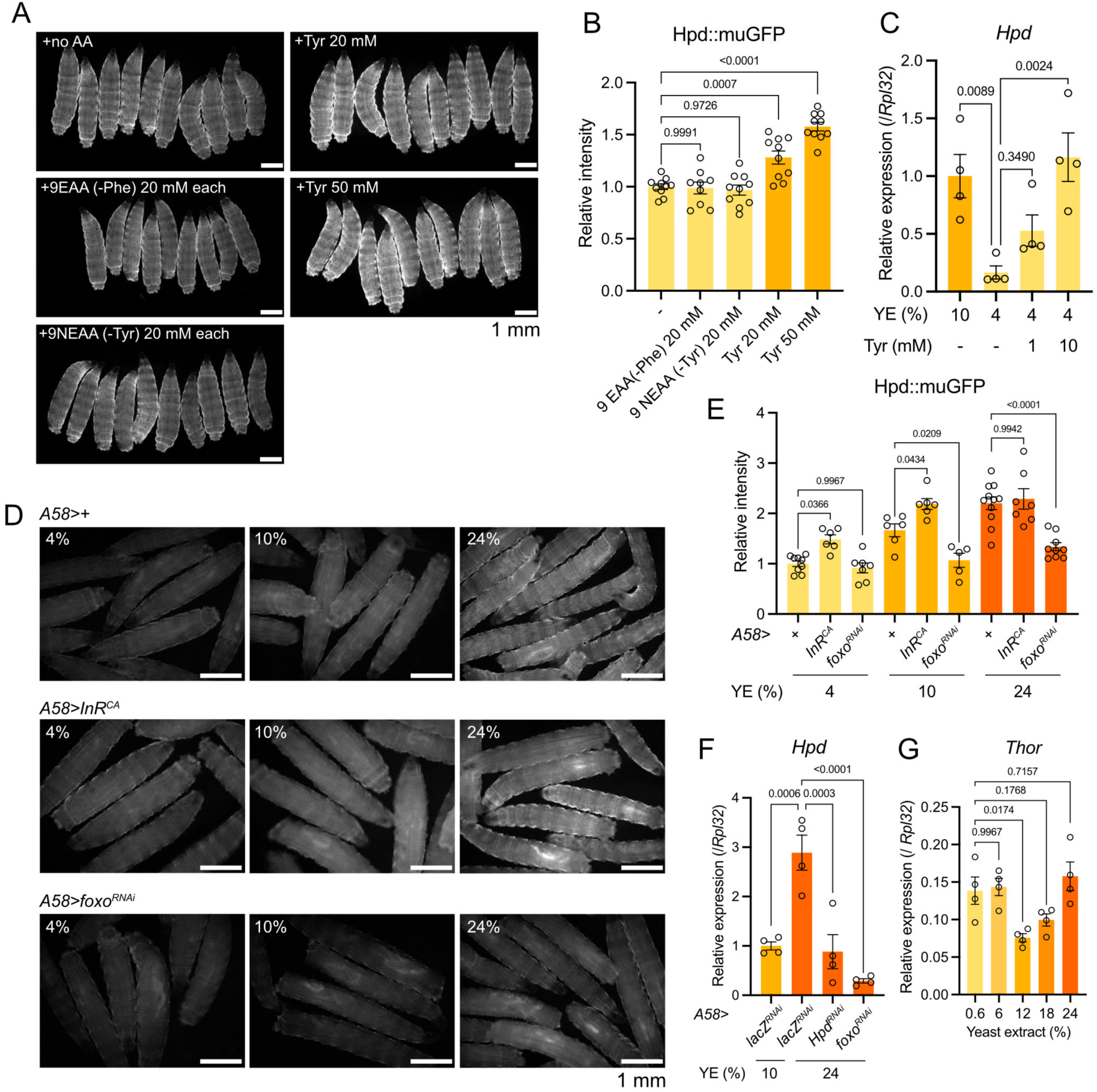
FoxO induces Hpd in response to high Tyr in the diet. (A, B) Images (A) and quantification (B) of *Hpd::muGFP* larvae upon AA supplementation to a standard yeast-based diet. Scale bar: 1 mm. n = 9-10. (C) Quantitative RT‒PCR of *Hpd* in the body wall of third instar larvae upon protein restriction with Tyr supplementation for 24 h. n = 4. (D, E) Images (D) and quantification (E) of *Hpd::muGFP* larvae upon various concentrations of dietary protein. Overexpression of *InR^CA^*or knockdown of *foxo* using epidermis driver (*A58-Gal4*). Scale bar: 1 mm. n = 5-11. (F) Quantitative RT‒PCR of *Hpd* in the body wall of third instar larvae upon protein restriction with Tyr supplementation for 24 h. Knockdown of *lacZ*, *Hpd*, or *foxo* using epidermis driver (*A58-Gal4*). n = 4. (G) Quantitative RT‒PCR of *Thor* in the whole body of third instar larvae upon different concentrations of dietary protein for eight hours. n = 4. For all graphs, the mean and SEM with all data points of biological replicates are shown. P-values were determined by one-way ANOVA with Dunnett’s multiple comparison test (B, G) and Holm-Šídák’s multiple comparison test (C, E, F).

It is rational to regulate the expression of the degradation enzyme by the excess amount of the substrate; however, the feedback mechanism for *Hpd* expression is not understood. Our previous study showed that the larval fat body senses Tyr scarcity via ATF4 (Kosakamoto et al., 2022). However, knockdown or overexpression of *ATF4* in the epidermis did not alter *Hpd* expression (Figs. S3B, C). In *C. elegans*, IIS induces Hpd protein expression via downregulation of DAF-16, an orthologue of FoxO (Lee et al., 2003). In rat hepatoma cells, the translation rate and the activity of the Tyr degradation enzyme Tat are increased by insulin treatment (Moore and Koontz, 1989). To test whether IIS is involved in Hpd regulation in *Drosophila*, we overexpressed the constitutively active form of insulin receptor (InR) in animals with *Hpd::muGFP*. Indeed, the activation of IIS mildly increased Hpd expression upon a low- or middle-protein diet (YE 4% or 10%) (Figs. 3D, E). However, overexpression of the dominant negative form of *insulin-like receptor* (*InR*) did not decrease Hpd expression upon a high-protein diet (Fig. S3B, C), suggesting that IIS is not the sole factor that regulates Hpd expression.

We next asked if FoxO is involved in Hpd regulation. Interestingly, Hpd expression upon a middle- or high-protein diet (YE 10% or 24%) was suppressed by *foxo*-RNAi (Figs. 3D, E). The transcription level of *Hpd* was also downregulated by *foxo*-RNAi, even more drastically than by *Hpd*-RNAi (Fig. 3F). This outcome was unexpected since FoxO was believed to be activated by protein restriction by IIS and therefore inactivated by IIS in a high protein context. To test whether FoxO is active in the epidermis of flies fed a high-protein diet, we performed qRT‒PCR of the FoxO target gene Thor/4E-BP. Intriguingly, *Thor* expression showed a U-shaped curve: it was induced upon both low- and high-protein diets (Fig. 3G). These results together suggest that Hpd expression is induced by FoxO when larvae are fed a high-protein diet, theoretically independent of IIS. In contrast, suppressed IIS in a low-protein diet leads to downregulation of Hpd expression in a FoxO-independent manner (Figs . 3D, E).

### Hpd induction is mediated by AMPK signalling

Our data suggested that high-protein stress activated FoxO by unknown mechanisms. To identify the upstream regulator of FoxO, we knocked down kinases that phosphorylate FoxO and promote its nuclear translocation. Jun amino terminal kinase (JNK) and AMP-activated protein kinase (AMPK) are two primary FoxO kinases (Brown and Webb, 2018). Overexpression of *puckered* (puc) or dominant negative form of *basket* (bsk) can suppress JNK signalling, but these manipulations did not suppress Hpd expression (Figs. S4A, B). In contrast, *AMPK* knockdown suppressed Hpd expression (Figs. 4A-C), suggesting that AMPK activates FoxO in the high protein diet and upregulates Hpd expression.

**Fig. 4.**
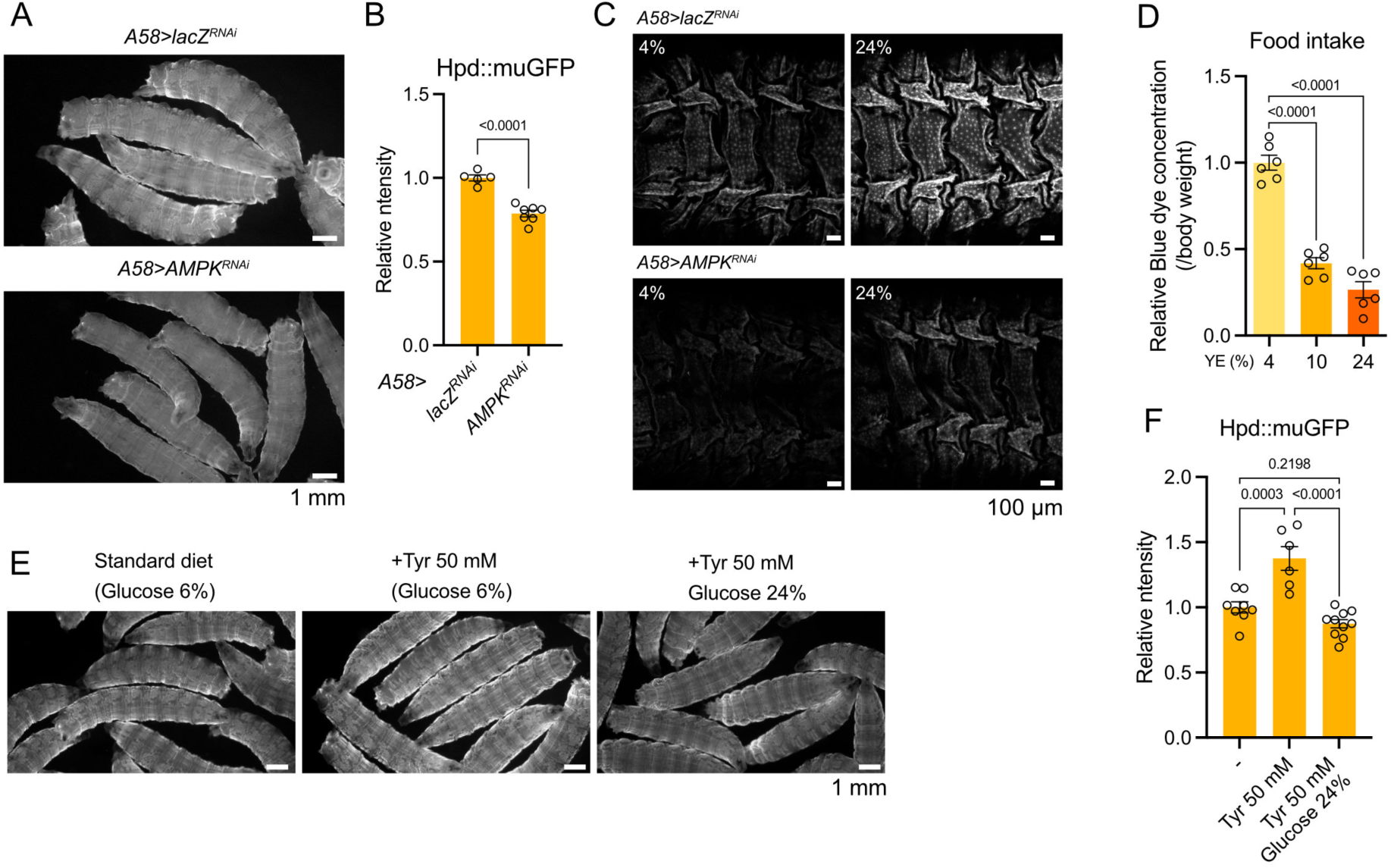
Hpd induction is regulated by AMPK signalling. (A, B) Images (A) and quantification (B) of *Hpd::muGFP* larvae fed the standard yeast-based diet. Knockdown of *AMPK* using epidermis driver (*A58-Gal4*). Scale bar: 1 mm. n = 5-7. (C) Images of dissected body walls of *Hpd::muGFP* larvae. Scale bar: 100 µm. (D) Food intake of wild-type larvae after being fed various concentrations of dietary protein for 24 h. n = 6. (E, F) Images (E) and quantification (F) of *Hpd::muGFP* larvae in the standard yeast-based diet with Tyr and glucose supplementation. Scale bar: 1 mm. n = 6-10. For all graphs, the mean and SEM with all data points of biological replicates are shown. P-values were determined by two-sided Student’s t-test (B), one-way ANOVA with Dunnett’s multiple comparison test (D) and Holm-Šídák’s multiple comparison test (F).

It is known that AMPK can recognize energy shortage by sensing the ATP/AMP ratio (Hardie, 2014). It is conceivable that the energy requirement is not met when larvae are fed a high-protein diet. Given that dietary protein is a critical regulator of food intake, we speculated that excess AA supply from a high YE diet suppresses food intake and therefore decreases carbohydrate intake. As expected, larvae fed a high-protein diet exhibited reduced food intake (Fig. 4D). Indeed, supplementation of glucose with a high Tyr diet suppressed Hpd expression (Figs. 4E, F). These results suggest that a high-protein diet activates the AMPK-FoxO axis partly due to insufficient sugar intake, which induces Hpd expression in the epidermis and maintains AA homeostasis.

### Hpd is essential for survival under protein overfeeding

To elucidate the physiological meaning of *Hpd* induction, we assessed how Hpd induction and accelerated Tyr metabolism contribute to the metabolic homeostasis and survival of animals. Notably, while control larvae exhibited the ability to maintain internal Tyr levels despite being fed a high-protein diet, Hpd knockdown in the epidermis led to a massive increase in overall Tyr levels only under this dietary condition (Fig. 5A). In addition, the downstream metabolite dopamine also exhibited elevated levels, while DOPA showed no change (Figs. 5B, C). Therefore, Hpd induction is necessary for larvae to maintain Tyr metabolism during protein overfeeding. As anticipated, *Hpd* knockdown, *Hpd* mutation, or *foxo* mutation all significantly impaired development when on high-protein diets, leading to lethality in both larval and pupal stages (Figs. 5D-I). These results emphasize the crucial role of Hpd in the epidermis for successful development under high-protein conditions (Fig. 5J).

**Fig. 5.**
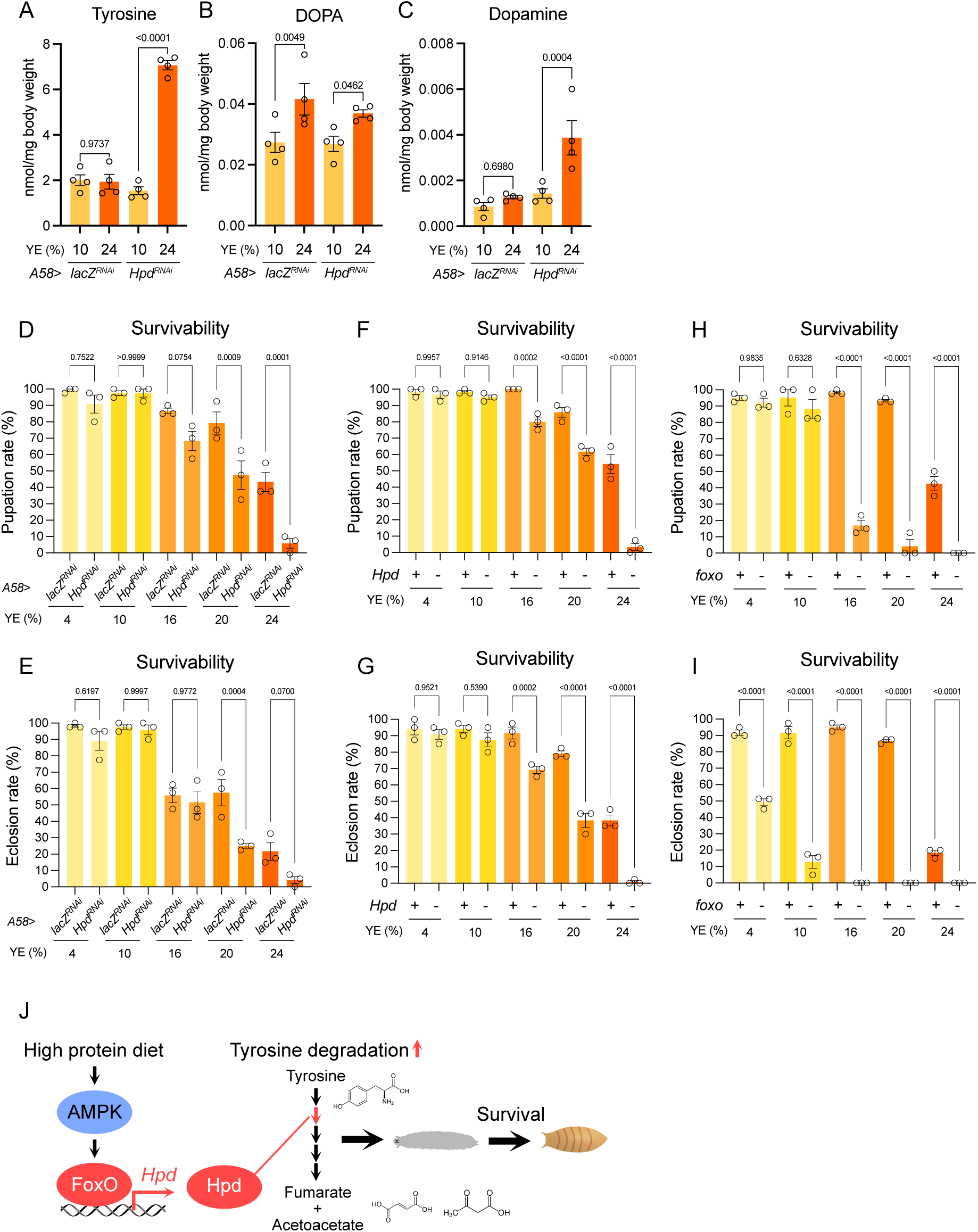
Hpd is necessary for Tyr homeostasis and survival upon protein overfeeding. (A-C) The internal level of Tyr metabolites in the larval whole body upon protein overfeeding for 24 h. Knockdown of *lacZ* or *Hpd* using epidermis driver (*A58-Gal4*). n = 4. (D-I) Pupation rate (D, F, H) or eclosion rate (E, G, I) of animals that have experienced various concentrations of dietary protein from third instar larvae. Knockdown of *lacZ* or *Hpd* using an epidermal driver (*A58-Gal4*) (D, E). Mutant of *Hpd* (F,G) or *foxo* (H,I). n = 3. For all graphs, the mean and SEM with all data points of biological replicates are shown. P- values were determined by one-way ANOVA with Holm-Šídák’s multiple comparison test. (J) The model of organismal adaptation to a high-protein diet.

## Discussion

In this study, we found the pivotal role of Tyr metabolism in epidermal cells for animal survival upon high-protein stress in *Drosophila* larvae. Intriguingly, the Tyr degradation pathway has also been shown to be crucial for the survival of hematophagous animals, which experience an increase in hazardous Tyr levels after blood digestion that must be mitigated through degradation (Sterkel et al., 2016). Hpd-silenced animals exhibit Tyr precipitation in the gut after a blood meal, which can result in lethality (Sterkel et al., 2016). Given that blood-sucking animals frequently transmit infectious diseases, vector control is of utmost importance, although the precise mechanism by which Tyr degradation enzymes are regulated *in vivo* is not yet entirely understood. This study reveals that the AMPK-FoxO axis regulates extensive Tyr degradation in the insect epidermis, providing valuable insights for efficient vector control in the future. Notably, Hpd inhibition did not kill *Drosophila* larvae fed a normal diet. Therefore, selective vector control, which is less damaging to other insects and ecosystems, could be achieved via the regulation of Tyr catabolism.

A previous study reported that the Tyr degradation pathway is related to lifespan regulation (Parkhitko et al., 2020). The extension of lifespan in long-lived flies is attributed to the downregulation of Tyr degradation and an increase in dopamine levels. In *C. elegans*, Hpd is an important target of DAF-16 (a FoxO orthologue) for lifespan extension in the *daf-2* (an orthologue of *InR*) mutant (Lee et al., 2003). The DAF16 and FoxO binding sites are conserved in the upstream region of the *Hpd* gene in both *C. elegans* and *D. melanogaster* (Lee et al., 2003). Our data showed that FoxO upregulates Hpd during high-protein feeding, whereas Lee et al. found that DAF16 downregulates Hpd. This contradiction may be due to different nutritional conditions, where excessive protein intake disrupts energy (sugar) homeostasis in our context. We found that AMPK is responsible for Hpd induction during high-protein stress. The ratio of dietary protein/carbohydrates determines larval food intake, and the reduced sugar intake activates AMPK. Interestingly, the elevation of internal Tyr levels by a genetic mutation in the *Tat* gene in *C. elegans* activates AMPK by unknown mechanisms (Ferguson et al., 2013). Furthermore, it has been shown that AMPK is activated by β-hydroxybutyrate derived from Tyr catabolism (Tong et al., 2021). These findings suggest that feed-forward regulation increases *Hpd* expression when Tyr levels are elevated. Since we found that Tyr specifically regulates Hpd expression among AAs, it is conceivable that AMPK is directly regulated by Tyr or its metabolite levels. A comprehensive examination of the mechanisms involved in AMPK-FoxO regulation in high-protein stress is needed.

In humans, Tyr catabolism is primarily regulated in the liver and to a lesser extent in the kidney (Noda and Ichihara, 1976). It has been reported that patients with tyrosinemia, a condition marked by a deficiency in the enzymes involved in Tyr degradation, suffer from tissue damage and disruption of hepatic and neurological functions due to crystal formation of Tyr and accumulation of intermediate metabolites from the Tyr degradation pathway (Adnan and Puranik, 2022; Najafi et al., 2018; Xie et al., 2019). To mitigate symptoms, these patients must block the formation of toxic Tyr metabolites and decrease dietary intake of Phe and Tyr (Adnan and Puranik, 2022), emphasizing the evolutionarily conserved significance of proper Tyr metabolism in humans. Recent studies identified Tyr as a predictive risk factor for the progression of diabetes and obesity (Newgard et al., 2009; Stancáková et al., 2012; Wang et al., 2011; Würtz et al., 2012). Additionally, decreased expression of Tat and Hpd in the liver is associated with poor prognosis in cancer patients, highlighting the role of Tyr metabolism in tumour suppression (Fu et al., 2010; Tong et al., 2021; Yang et al., 2020). Therefore, a more profound comprehension of the regulatory mechanisms governing Tyr metabolism has the potential to inform effective clinical interventions for such diseases.

The *Drosophila* counterparts of the mammalian liver are the fat body and oenocyte (Gutierrez et al., 2007; Li et al., 2019). While the fat body serves as a primary metabolic organ, it appears to not participate in Tyr degradation. Our previous research has shown that the fat body is an essential organ for sensing Tyr to adapt to a dietary protein environment (Kosakamoto et al., 2022). Thus, interorgan communication likely exists between the epidermis, which maintains internal Tyr levels, and the fat body, which monitors Tyr levels. The specialized role of the epidermis in Tyr degradation may be attributed to the organ’s significant demand for Tyr during metamorphosis for pigmentation and sclerotization (Gorman and Arakane, 2010). It is reasonable for *Drosophila* larvae to make use of those enzymes already existing in the epidermis to counteract high protein toxicity rather than expressing them in the fat body. High levels of Tyr are stored in vacuoles within the insect fat body and released into the haemolymph during metamorphosis (McDermid and Locke, 1983). The transport of large amounts of Tyr and acute promotion of melanogenesis carry a risk of oxidative damage. Therefore, it is likely that the degradation machinery for Tyr is necessary in the epidermis to prevent damage to the organ. Nonetheless, even after completing metamorphosis, adult flies also show Hpd expression in epidermal tissue as well as oenocytes. The evolutionary specialization of the epidermis in Tyr degradation in insects is a fascinating question. Together, the unexpected participation of the structural component in animal metabolic homeostasis would broaden the perspective of the adaptation mechanism to a changing nutritional environment.

## Materials and Methods

### *Drosophila* stocks and husbandry

Flies were reared on a standard yeast-based diet containing 4.5% cornmeal (NIPPN CORPORATION), 6% brewer’s yeast (ASAHI BREWERIES, HB-P02), 6% glucose (Nihon Shokuhin Kako), and 0.8% agar (Ina Food Industry S-6) with 0.4% propionic acid (Wako 163-04726) and 0.15% butyl p-hydroxybenzoate (Wako 028-03685). Flies were maintained under conditions of 25°C. To allow synchronized development and constant density, embryos were collected by agar plates (2.3% agar, 1% sucrose, and 0.35% acetic acid) with live yeast paste.

The fly lines used in this study were *w^Dah^* (Grandison et al., 2009), *foxo^Δ94^* (Slack et al., 2011), *A58-Gal4* (Galko and Krasnow, 2004), *UAS-mCD8-RFP* (Bloomington Drosophila Stock Center (BDSC) 27391), *UAS-GFP* (BDSC 6874), *UAS-ATF4* (FlyORF, F000106), *UAS-Hpd-RNAi* (Vienna Drosophila Resource Center (VDRC) 103482), *UAS-foxo-RNAi* (BDSC 32429), *ATF4-RNAi* (VDRC 109014), *UAS-lacZ-RNAi, UAS-InR^DN^* (BDSC 8253), *UAS-InR^CA^* (BDSC 8263), *UAS- UAS-bsk^DN^* (Kyoto Drosophila Stock Center, 108773), and *UAS-puc* (from Dr. Igaki).

To generate *Hpd::muGFP* flies, we used the CRISPR/Cas9 system to insert *monomeric ultrastable GFP* (*muGFP*) at the C-terminus of the *Hpd* gene (Kina et al., 2019). The codons of *muGFP* were optimized for expression in *Drosophila melanogaster*. These modifications and insertions into the EcoRI/XbaI site in the pUC57 vector were performed by GenScript. An sgRNA target site of *Hpd* was selected using CRISPR Optimal Target Finder (Gratz et al., 2014). Complementary oligonucleotides with overhangs were annealed and cloned into the BbsI-digested U6b vector using a DNA ligation kit (Takara, 6023).

Sense strand: TTCGGAAATTGAACAAGCCAAGCG

Antisense strand: AAACCGCTTGGCTTGTTCAATTTC

The targeting vector was constructed by inserting *muGFP* between 500 bp of *Hpd* homology arms with a mutation in the PAM sequence next to the sgRNA site (from AGG to AGA). First, *muGFP* and homology arms were PCR-amplified using Q5 High-Fidelity 2× Master Mix (New England BioLabs, M0492L). Primers for PCR were designed using the NEBuilder Assembly Tool. The gel-purified PCR products were cloned into the EcoRI-digested pBluescript II SK(+) vector using NEBuilder HiFi DNA Assembly Master Mix (New England BioLabs, E2621X). The mixture of pU6b-sgRNA and targeting vector was microinjected into w1118; attP40{nos-Cas9}/CyO embryos by WELLGENETICS. The F0 adults were crossed with balancer lines, and the *muGFP*-inserted lines were selected by PCR amplification of the target locus. The primers used for PCR are listed in Table S1.

To generate *pHpd-GeneSwitch* flies, 500 bp upstream of the start codon of *Hpd* and the *GeneSwitch* sequence were PCR-amplified using Q5 High-Fidelity 2× Master Mix.

Primers for the PCR were designed using the NEBuilder Assembly Tool. The gel-purified PCR products were cloned into the KpnI-digested pElav-GeneSwitch vector using NEBuilder HiFi DNA Assembly Master Mix. The plasmid was injected into w1118 embryos by WELLGENETICS. The F0 adults were crossed with balancer lines, and w+ lines were selected. The primers used for PCR are listed in Table S1.

### Dietary manipulations

Embryos were harvested under the standard yeast-based diet until they became third instar larvae, especially 72 h after egg laying (AEL), unless otherwise stated. Then, the larvae were floated using 30% glycerol and transferred through a soft brush to fly vials containing various diets. For protein manipulation, 4%, 10%, or 24% yeast extract (Nacalai, 15838-45) was mixed with 6% glucose, 1% agar, 0.3% propionic acid and 0.15% nipagin (YE diet).

For the survival assay, we added 1% baker’s yeast (LESSAFLE, Saf-instant yeast red) to the YE diet above to promote larval development.

### RNA sequencing analysis and quantitative RT‒PCR

For RNAseq analysis of larval carcasses, the larvae were fed three types of diets for six hours from 90 h AEL. Total RNA was purified from six carcasses of female third instar larvae using a Promega ReliaPrep RNA Tissue Miniprep kit (z6112). Triplicate samples were prepared for each experimental group. RNA was sent to Kazusa Genome Technologies to perform 3’ RNA-seq analysis. The cDNA library was prepared using the QuantSeq 3’ mRNA-Seq Library Prep Kit for Illumina (FWD) (LEXOGEN, 015.384).

Sequencing was performed using Illumina NextSeq 500 and NextSeq 500/550 High Output Kit v2.5 (75 cycles) (Illumina, 20024906). Raw reads were analysed by the BlueBee Platform (LEXOGEN) for trimming, alignment to the *Drosophila* genome, and counting of the reads. The count data were analysed by the Wald test using DESeq2. The result has been deposited in DDBJ under accession number DRA016117.

For quantitative RT‒PCR analysis, total RNA was purified from the whole body or dissected carcass from third instar larvae using a Promega ReliaPrep RNA Tissue Miniprep kit (z6112). The cDNA was synthesized from 400 ng of DNase-treated total RNA by the Takara PrimeScript RT Reagent Kit with gDNA Eraser (Takara bio RR047B). Quantitative PCR was performed using TB Green™ Premix Ex Taq™ (Tli RNaseH Plus) (Takara bio RR820W) and a QuantStudio 6 Flex Real Time PCR system (ThermoFisher) using *Rpl32* as an internal control. Primer sequences are listed in Supplementary Table 1.

### Measurement of metabolites

Metabolites were measured by ultra-performance liquid chromatography-tandem mass spectrometry (LC‒MS-8050/LCMS-8060, Shimadzu) based on the Primary metabolites package ver.2 (Shimadzu) (Kosakamoto et al., 2022; Shiota et al., 2018). Five whole bodies of third instar larvae were homogenized in 160 μL of 80% methanol containing 10 μM internal standards (methionine sulfone and 2-morpholinoethanesulfonic acid). After centrifugation at 14,000 × g and 4°C for five minutes, 150 μL of supernatant was mixed with 75 μL of acetonitrile and deproteinized. After centrifugation at 20,000 × g and 4°C for five minutes, the supernatant was applied into a prewashed 10-kDa centrifugal device (Pall, OD010C35), and the flow-through was completely evaporated using a centrifugal concentrator (TOMY, CC-105). The samples were resolubilized in ultrapure water and injected into the LC‒MS/MS with a PFPP column (Discovery HS F5 (2.1 mm × 150 mm, 3 μm), Sigma‒Aldrich) in the column oven at 40°C. A gradient from solvent A (0.1% formic acid, water) to solvent B (0.1% formic acid, acetonitrile) for 20 min was used to separate solutes. The MRM parameters were optimized by the injection of the standard solution through peak integration and parameter optimization with the software (Labsolutions, Shimadzu).

### Imaging analysis

To analyse Hpd::muGFP expression in larvae, the animals were picked up and left in embryo dishes filled with Milli-Q water for eight minutes at -30°C. When larval movement was restricted, images were acquired using a fluorescence stereomicroscope (Leica MZ10F). GFP fluorescence was quantified using Fiji by calculating the average intensity of three regions of interest between body segments of each larva.

For more detailed analysis of the expression pattern of Hpd, the larval epidermis was dissected as described previously (Ramachandran and Budnik, 2010). Briefly, a larva was picked up and placed in a drop of PBS and then pinned at the anterior and posterior ends with two pins. To observe the dorsal side, the ventral side is placed at the top. Using micro scissors, cut the larva from the posterior side to the anterior end. After removing the internal organs, the epidermis was gently stretched and pinned by four corner pins. The sample was fixed by replacing PBS with 4% paraformaldehyde (PFA) in PBS. After fixation for 20 min at room temperature (RT), the sample was washed multiple times with PBST (0.1% TritonX-100) and incubated for 2 h at RT with Hoechst 33342 (Invitrogen, H3570, diluted to 0.4 mM) at 1:100 and Phalloidin-iFluor 647 (Abcam, ab176759, 1:5000). After washing, tissues were mounted in 80% glycerol and observed using a Leica TCS SP8 confocal microscope.

### Feeding assay

Embryos were collected within 4 h of egg laying using a small cage and harvested in a bottle containing a standard diet (13 μL/bottle). At 60 h after egg laying, second instar larvae were floated up using 30% glycerol and gently transferred to each diet (YE4%, 10%, and 24%). After 24 h, the third instar larvae were again floated up and gently transferred to a drop of each diet containing 1% blue dye (Wako, 027-12842) on the centre of a 35 mm plate. The diet was softened in advance by scratching the surface with a spatula to make it easy for larvae to crawl into the food. After 15 min of feeding, the larvae were floated with 30% glycerol, and the blue dye on the body wall was extensively washed with Milli-Q water. The body surface was swiftly wiped with Kimwipes, and eight larvae were collected in a 1.5-mL tube containing 80 μL of 100% methanol. Larvae were homogenized using a pellet pestle and centrifuged for 5 min at 20,000 ×g. The supernatant was used for measuring A620 to quantify the blue dye concentration (InfiniteF50R, Tecan).

### Statistical analysis

Statistical analysis was performed using GraphPad Prism 8 or 9. The sample numbers were determined empirically. All data points were biological, not technical, replicates. No data were excluded. An unpaired and two-sided Student’s *t*-test was used to test between samples. One-way ANOVA with Holm-Šídák’s multiple comparison test was used to test among groups. One-way ANOVA with Dunnett’s multiple comparison test was used to test with a control sample. All experimental results were repeated at least twice to confirm the reproducibility. Bar graphs are drawn as the mean and SEM.

## Acknowledgments

We would like to acknowledge Kyoto Stock Center, National Institute of Genetics, Vienna Drosophila Resource Center, and Bloomington Drosophila Stock Center for reagents. We thank all members of our laboratory for technical assistance and critical advice.

## Author Contributions

H.K. and F.O. conceived the project and wrote the manuscript. H.K. performed most of the experiments and analyzed the data. M.M. and F.O. supervised the study. All authors edited and approved the final manuscript.

## Data availability

The NGS data are available under accession numbers, DRA016117. The data used to analyze the results in this paper are available as source data. All the materials generated in this study are available upon request to F.O.

---

**Fig. S1.**
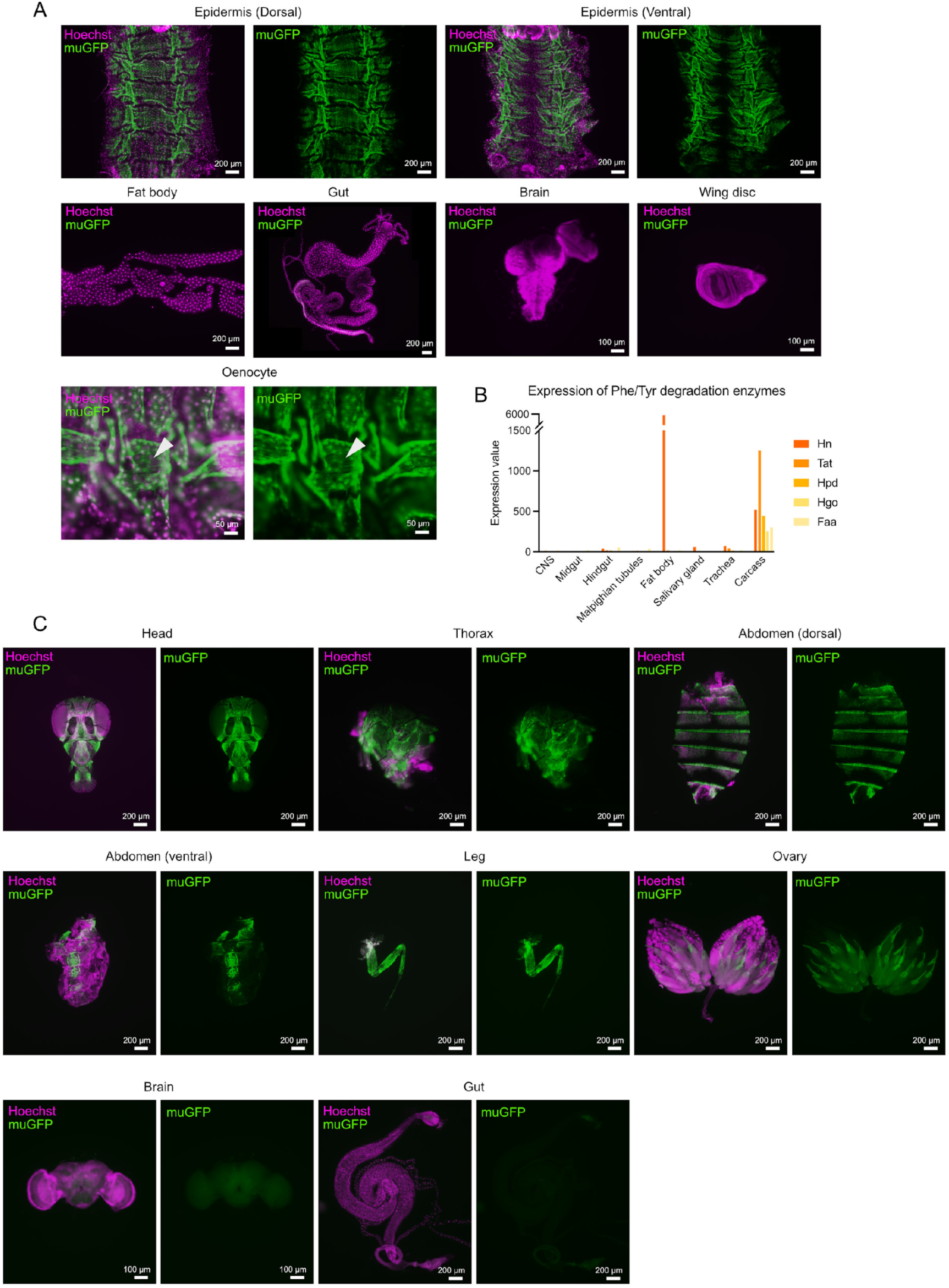
Expression of Tyr degradation enzymes are enriched in the carcass. (A) Images of several tissues of *Hpd::muGFP* larvae upon a high-protein diet (24% YE) for 24 hours. (B) Expression levels of Phe and Tyr degradation enzymes from microarray dataset of flybase. (C) Images of several tissues of *Hpd::muGFP* adults upon a high-protein diet (24% YE) for four days.

**Fig. S2.**
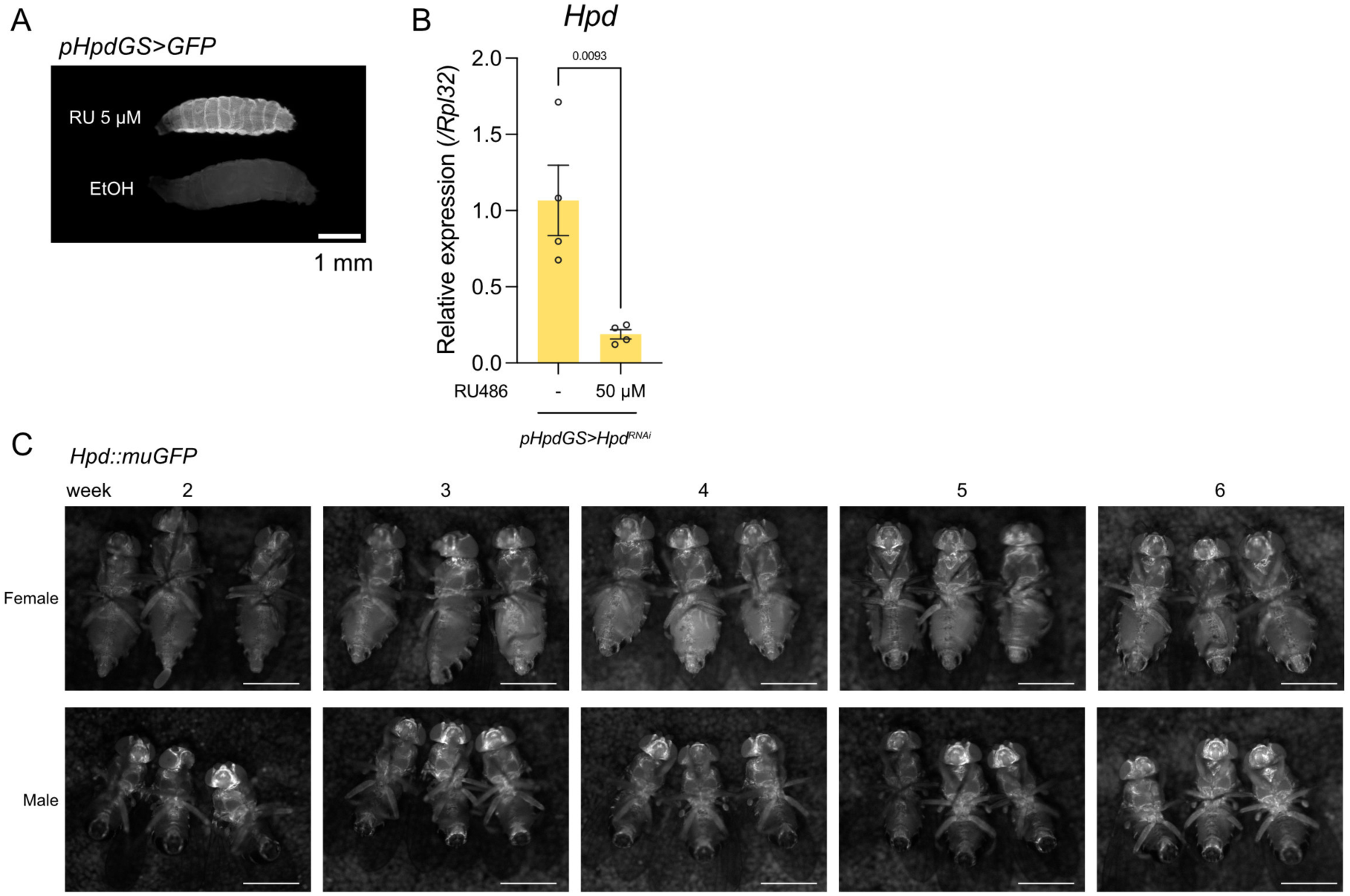
Generation of epidermis GeneSwitch driver and changes of Hpd expression during aging. (A) A representative image of *pHpdGS>GFP* larvae with or without mifepristone (RU486) feeding. scalr bar: 1 mm. (B) Quantitative RT-PCR of *Hpd* in the body wall of third instar larvae upon a high-protein diet (24%YE) with RU486 for 24 hours. Knockdown of *Hpd* using *pHpdGS*. n = 4. (C) Images of *Hpd::muGFP* adults upon standard yeast based diet during aging. Scale bar: 1 mm. For the graph, mean and SEM with all data points of biological replicates were shown. P-values are determined by two-sided Student’s t-test.

**Fig. S3.**
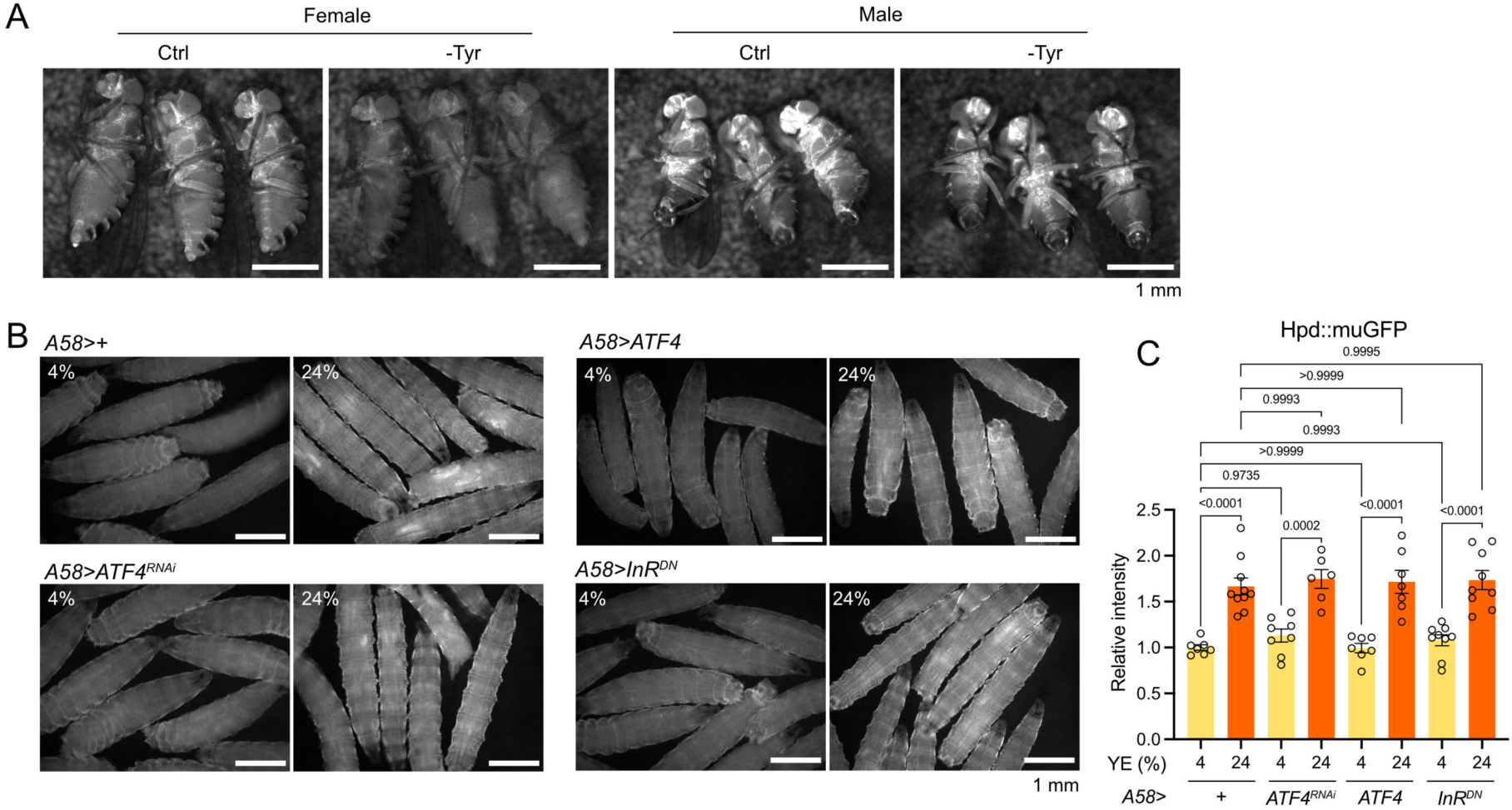
Tyr, but not ATF4 signaling, is involved in Hpd regulation. (A) Images of *Hpd::muGFP* adults upon synthetic diet with or without Tyr for three days from five days after eclosion. Scale bar: 1 mm. (B,C) Images and the quantification (C) of *Hpd::muGFP* larvae upon a low- or high-protein diet. Knockdown or overexpression of *ATF4* or *InR^DN^* using epidermis driver (*A58-Gal4*). Scale bar: 1 mm. n = 6-10. For the graph, mean and SEM with all data points of biological replicates were shown. P-values are determined by one-way ANOVA with Holm-Šídák’s multiple comparison test.

**Fig. S4.**
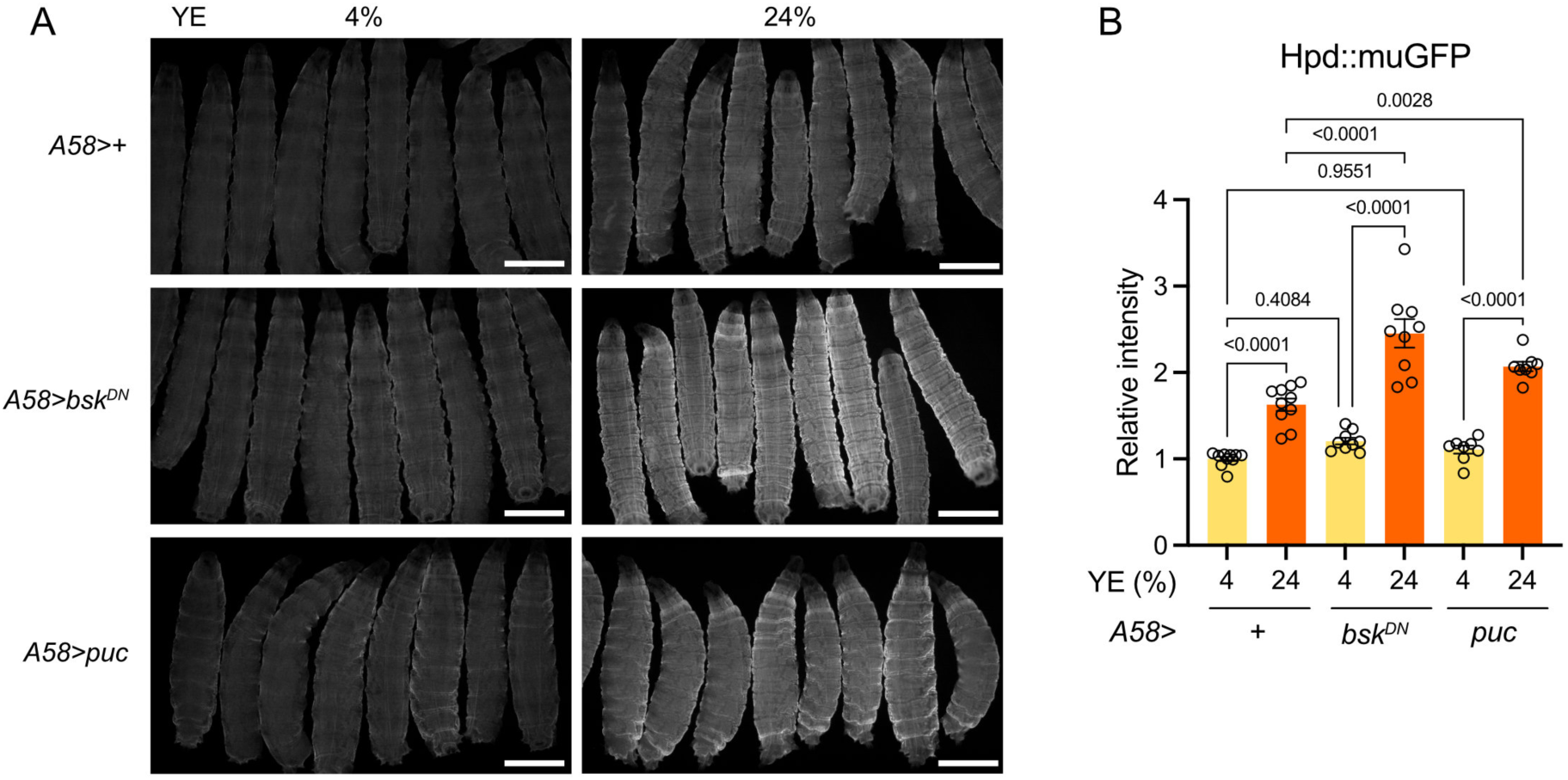
JNK signaling is not involved in Hpd signaling. (A,B) Images and the quantification (B) of *Hpd::muGFP* larvae upon a low- or high-protein diet. Overexpression of *bsk^DN^* or *puc* using epidermis driver (*A58-Gal*4). Scale bar: 1 mm. n = 8-10. For the graph, mean and SEM with all data points of biological replicates were shown. P-values are determined by one-way ANOVA with Holm-Šídák’s multiple comparison test.

**Table S1.**
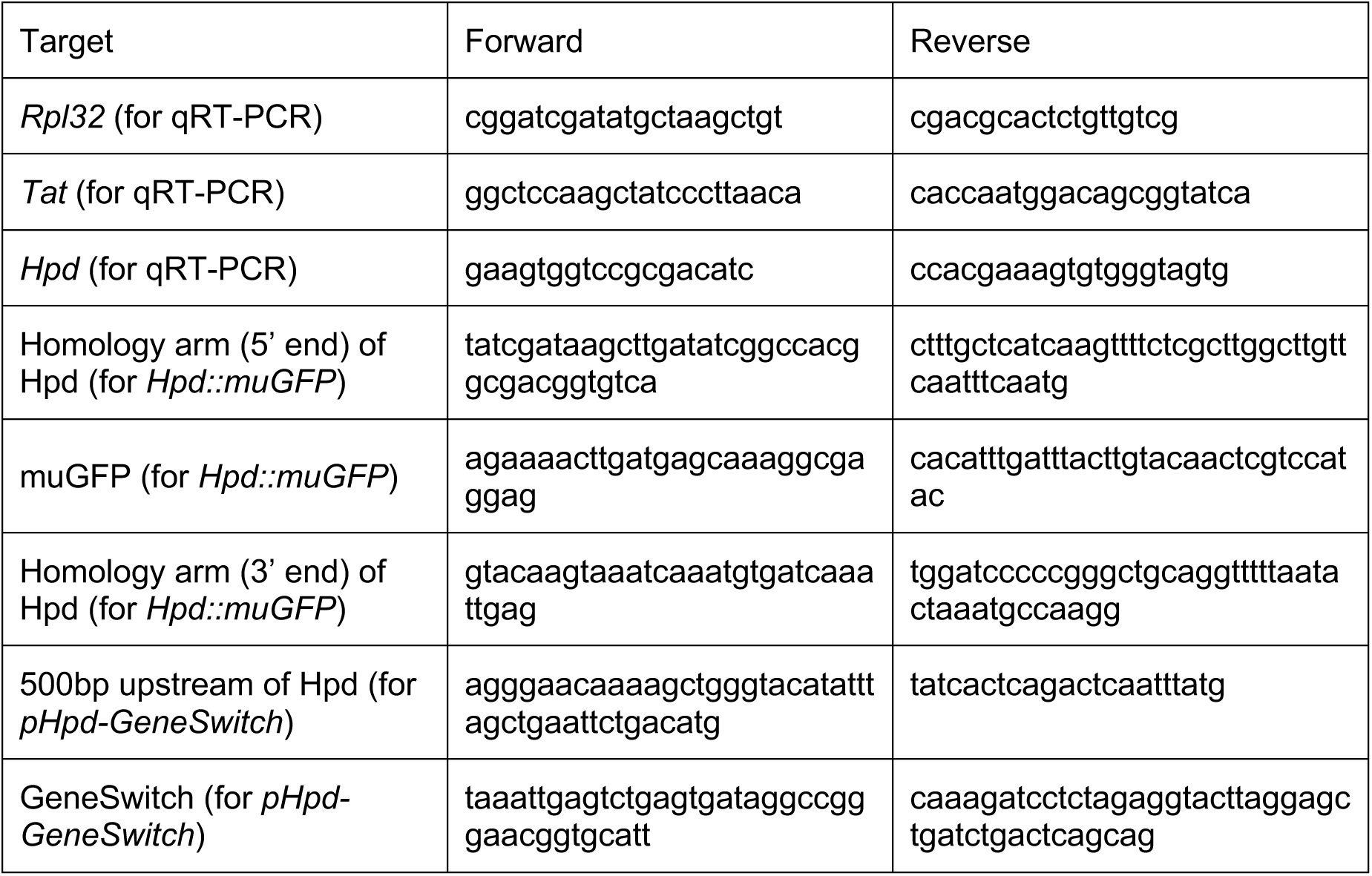
Primers used in this study.

